# TOXsiRNA: A web server to predict the toxicity of chemically modified siRNAs

**DOI:** 10.64898/2026.02.12.705521

**Authors:** Showkat Ahmad Dar, Manoj Kumar

## Abstract

Small interfering RNAs (siRNAs) are largely modified with chemical molecules to enhance their properties for use in molecular biology research and therapeutic applications. Toxicity effects may arise due to these chemical moieties as well as sequence based off-targets at cellular level. Enormous resources are required to experimentally design and test the toxicity of these chemical modifications and their combinations on siRNAs. To address this problem, we developed TOXsiRNA web server to computationally predict the toxicity of chemically modified siRNAs and their off-targets. We selected 2749 siRNAs with different permutations and combinations of 21 different chemical modifications engineered on them. Next, we used Support Vector Machine (SVM), Linear Regression (LR), K-Nearest Neighbor (KNN) and Artificial Neural Network (ANN) machine learning applications to develop models. Best performance was displayed by mononucleotide composition-based model developed with SVM, offering Pearson Correlation Coefficient (PCC) of 0.91 and 0.92 on training testing and independent validations respectively. Other sequence features like dinucleotide composition binary pattern and their combinations were also tested. Finally, three models of chemically modified siRNAs were implemented on the web server. Other algorithms that include predicting normal as well as chemically modified siRNA knockdown efficacy, off target etc. are also integrated. The resource is hosted online for scientific use freely at url: http://bioinfo.imtech.res.in/manojk/toxsirna.

## Introduction

The siRNAs are employed for gene knockdown in research and tried as potential therapeutics for various diseases ranging from cancers, viral infections, metabolic diseases to neurological abnormalities etc. [1-3]. Though, certain inherent physicochemical properties limit the use of these natural siRNAs (unmodified) as therapeutic agents [4-6]. To overcome these shortcomings, they are modified with different or same kind of chemical molecules with different permutations and combinations on either or both strands [4]. These modifications improved the performance of the siRNAs e.g. the knockdown efficacy of siRNA was reported to be boosted 500 times with respect to its unmodified counterpart [7]. Similarly, many other such reformations in the normal siRNAs have lead to development of second generation of siRNAs (chemically modified siRNAs), which are very promising in devising therapeutic deliverables [2]. All the constituent moieties of nucleotide (ribose sugar, nitrogenous base and the phosphate group) are modified and the ribose predominates statistically [4]. Recently, siRNA based drug molecule Patisiran (Onpattro^™^), a 2’-O-methyl chemically modified is approved by the FDA for clinical applications against the hereditary transthyretin-mediated amyloidosis (hATTR) [8, 9].

Among the various limitations of the siRNAs that needs to be taken care of, toxicity is one of the important aspects that arise either due to off targets or due to chemical modifications [10, 11]. Off target based toxicity may arise from base pairing of the siRNA to any other mRNAs sequences that have potential base pairing probability (phenomena similar to that of miRNA) [10, 12, 13]. Nonetheless, these additional target bindings are suppressed with certain chemical modifications [10, 14]. For an instance, the second nucleotide antisense strand when modified with 2’-O-Methyl ribose chemical moiety displays reduced off target effects [14]. Similarly, the chemical modification with locked nucleic acid demonstrated very few off target effects against its counterpart [15]. Conversely, an advantage of the off target effects can be applied in the process of “death induced by survival gene elimination” (DISE) in the malignant cells to limit their growth [12] [16].

In literature, many computational algorithms exist that can design and determine the efficacy of the siRNAs (normal as well as chemically modified) but none provides information about the toxicity of the siRNAs [17, 18]. Furthermore, many computational methodologies have been reported in the literature that emphasize and show that such systems are needed for prediction of toxicity of different molecules [19, 20]. Thus, to predict the toxicity of the chemically modified siRNAs and their off-targets and provide the missing link in the field we constructed the TOXsiRNA web server.

## Results

### Machine Learning Model development

Mononucleotide composition (MNC) and Binary pattern (BIN) features with two types of arrangement of nucleotide sequence data resulting in six categories (three random and three sequential) for cross validation were used [18]. The performance of 10-fold cross validation for machine learning model with four techniques (LR, KNN, ANN and SVM) is shown in the **Table 1**. More or less all the four machine-learning techniques behaved equally for a particular feature on the dataset in terms of Pearson correlation coefficient (PCC). The **Table 1** revealed that for both sequential and random datasets, there is no significant difference in the performance of model development and both achieved almost equal PCC. Their corresponding PCC values (10-fold cross validations) were almost equal with minor variations in some of the methods. Subsequently, we opted the 5^th^ sequential sample dataset for model development for all features and with all the four machine-learning techniques. Simultaneously, various hybrid models for MNC and BIN features were used for model development. The result for multiple features and their combinations is shown in the **Table 2** with their corresponding PCC values of respective techniques.

**Table 1.** Sequential dataset (2^nd^, 5^th^ and 8^th^) and random datasets (R-I, R-II and R-III) three datasets each for mononucleotide composition (MNC) and binary models (BIN), their number of sequence features used for selecting the datasets for cross validation along with their corresponding Pearson Correlation coefficients (PCC).

|  | Model | Vector size | Data-set | LR | ANN | KNN | SVM |
| --- | --- | --- | --- | --- | --- | --- | --- |
|  |  |  | 10-Fold | PCC | PCC | PCC | PCC |
| 1 | MNC | 52 | 2nd | 0.84 | 0.85 | 0.83 | 0.91 |
| 2 | MNC | 52 | 5th | 0.84 | 0.83 | 0.83 | 0.91 |
| 3 | MNC | 52 | 8th | 0.83 | 0.83 | 0.83 | 0.91 |
| 4 | MNC | 52 | R-I | 0.83 | 0.85 | 0.83 | 0.91 |
| 5 | MNC | 52 | R-II | 0.82 | 0.82 | 0.84 | 0.91 |
| 6 | MNC | 52 | R-III | 0.82 | 0.83 | 0.83 | 0.91 |
| 7 | BIN | 48 | 2nd | 0.58 | 0.76 | 0.80 | 0.34 |
| 8 | BIN | 48 | 5th | 0.59 | 0.79 | 0.80 | 0.34 |
| 9 | BIN | 48 | 8th | 0.02 | 0.74 | 0.81 | 0.34 |
| 10 | BIN | 48 | R-I | 0.60 | 0.70 | 0.80 | 0.34 |
| 11 | BIN | 48 | R-II | 0.60 | -0.02 | 0.80 | 0.34 |
| 12 | BIN | 48 | R-III | 0.60 | 0.78 | 0.81 | 0.34 |

PCC =Pearson Correlation coefficient; SVM = Support Vector Machines; LR = Linear Regression; KNN = k-Nearest Neighbors; ANN = Artificial Neural Networks

**Table 2.** Single feature (1,2,3) and hybrid feature (4 to 15) models used for model development with four Machine-learning techniques using 5^th^ dataset. Sequence features, Vector size, the dataset used and PCC values of corresponding algorithms.

| S. No. | Feature used | Vector size | LR | ANN | KNN | SVM |
| --- | --- | --- | --- | --- | --- | --- |
|  |  |  | PCC | PCC | PCC | PCC |
| 1 | Mononucleotide (A) | 52 | 0.84 | 0.83 | 0.83 | 0.91<sup>#</sup> |
| 2 | Dinucleotide | 1352 | 0.86 | 0.85 | 0.15 | 0.88 |
| 3 | Binary (B) | 48 | 0.59 | 0.79 | 0.80 | 0.34 |
| 4 | (A+B) 24-ntd both strands | 100 | 0.78 | 0.81 | 0.83 | 0.33 |
| 5 | (A+B) 16-ntd both strands | 84 | 0.89 | 0.80 | 0.82 | 0.61 |
| 6 | (A+B) 13ntd both strands | 78 | 0.79 | 0.81 | 0.82 | 0.62 |
| 7 | (A+B) 8-ntd both strands | 68 | 0.82 | 0.82 | 0.82 | 0.71 |
| 8 | A+B 24-ntd Antisense only | 76 | 0.86 | 0.83 | 0.83 | 0.86<sup>#</sup> |
| 9 | A+B 16-ntd Antisense only | 68 | 0.80 | 0.83 | 0.83 | 0.88 |
| 10 | A+B 13ntd Antisense only | 65 | 0.80 | 0.79 | 0.83 | 0.88 |
| 11 | A+B 8-ntd Antisense only | 60 | 0.82 | 0.82 | 0.83 | 0.89 |
| 12 | A+B 24-ntd Sense only | 76 | 0.86 | 0.78 | 0.82 | 0.88<sup>#</sup> |
| 13 | A+B 16-ntd Sense only | 68 | 0.88 | 0.80 | 0.81 | 0.89 |
| 14 | A+B 13ntd Sense only | 65 | 0.83 | 0.80 | 0.82 | 0.89 |
| 15 | A+B 8-ntd Sense only | 60 | 0.83 | 0.80 | 0.82 | 0.90 |

PCC = Pearsons Correlation coefficient; SVM = Support Vector Machines; LR = Linear Regression; KNN = k-Nearest Neighbors; ANN = Artificial Neural Networks Final three models (#) selected from SVM machine learning technique for web server implementation.

### Independent validation

The independent validation of the two features MNC and BIN for each six datasets (three respectively for random and sequential) performed almost equally well as seen from their PCC values **Table 3**. Subsequently, we selected the three best models from all the models created, based on SVM technique as its performance was better. Their independent validation with IV-275 dataset also performed well in terms of PCC as seen in **Table 4**.

**Table 3.** The independent validation SVM model results for testing the Independent validation dataset of 275 sequences (IV-275) with corresponding PCC.

|  | **Models** | **Vector size** | **Datasets** | **PCC** | **TESTSET** |
| --- | --- | --- | --- | --- | --- |
| 1 | MNC | 52 | 2nd | **0.91** | 2474 |
|  |  | 52 | I-V-2 | **0.91** | 275 |
| 2 | MNC | 52 | 5th | **0.91** | 2474 |
|  |  | 52 | I-V-5 | **0.92** | 275 |
| 3 | MNC | 52 | 8th | **0.91** | 2474 |
|  |  | 52 | I-V-8 | **0.92** | 275 |
| 4 | MNC | 52 | R-I | **0.91** | 2474 |
|  |  | 52 | I-V-I | **0.92** | 275 |
| 5 | MNC | 52 | R-II | **0.91** | 2474 |
|  |  | 52 | I-V-II | **0.92** | 275 |
| 6 | MNC | 52 | R-III | **0.91** | 2474 |
|  |  | 52 | I-V-II | **0.91** | 275 |
| 7 | BIN | 48 | 2nd | **0.34** | 2474 |
|  |  | 48 | I-V-2 | **0.42** | 275 |
| 8 | BIN | 48 | 5th | **0.34** | 2474 |
|  |  | 48 | I-V-5 | **0.38** | 275 |
| 9 | BIN | 48 | 8th | **0.34** | 2474 |
|  |  | 48 | I-V-8 | **0.34** | 275 |
| 10 | BIN | 48 | R-I | **0.34** | 2474 |
|  |  | 48 | I-V-I | **0.36** | 275 |
| 11 | BIN | 48 | R-II | **0.34** | 2474 |
|  |  | 48 | I-V-II | **0.41** | 275 |
| 12 | BIN | 48 | R-III | **0.34** | 2474 |
|  |  | 48 | I-V-III | **0.36** | 275 |

R = random datasets; I-V = Independent validation dataset

**Table 4.** SVM models (with 10-fold and Independent validation datasets) with independent validation results that were selected for web server implementation conclusively.

|  | **Model** | **RUN** | **PCC** | **Dataset** |
| --- | --- | --- | --- | --- |
| I | MNC | 10-fold | **0.91** | 2474 |
|  |  | I-V | **0.92** | 275 |
| II | MNC+ BIN-24AS | 10-fold | **0.86** | 2474 |
|  |  | I-V | **0.83** | 275 |
| III | MNC+ BIN-24SS | 10-fold | **0.88** | 2474 |
|  |  | I-V | **0.89** | 275 |

Here: MNC = Mononucleotide composition; MNC+BIN-24SS =Mononucleotide and binary combined for Sense strand; and MNC+BIN-24AS =Mononucleotide and binary combined for Antisense strand; I-V = Independent validation dataset

### Web server components

Next, we applied the SVM based models developed along with additional programmes on the web-server that are listed below:

### TOXsiRNA

The TOXsiRNA option is the first part of the web server and can predict the normal siRNA (as well as chemically modified siRNAs but with limited modifications per sequence). The user can paste or upload the target sequence against which the siRNAs needs to be designed and upon submitting the job, it will produce the list of siRNAs with respective knockdown efficacy along with its off targets. These siRNAs are based on the SVM prediction model from siRNApred algorithm [18]. Each siRNA sequence from this list (sortable in ascending or descending order according to knockdown efficacy) can be clicked and will lead to next page with 882 (21 x 21 x 2), variants of chemically modified sequences. The chemical modifications are based on 21 chemical moieties at 21 position of either strand represented with the single letter code as shown in **Supplementary Table 1**. The 882 modified siRNAs are listed with their corresponding toxicity score ranging from 0 to 1 with the former being least toxic and higher value as most toxic. Additionally, the efficacy values are provided that are based our earlier work SMEpred [18].

### GENMod

AS we observed above, only single modification at single positions are generated. However, if the user wants to create the siRNA with multiple modifications and multiple positions and check its toxicity, GENMod programme will help. It provides the option to upload the normal nucleotide sequence with sense and antisense strands separated with hyphen (-) (e.g. GCAGCACGACUUCUUCAAGUUCUUGAAGAAGUCGUGCUGCUU). Next option is for chemical modifications separated by double commas (,,) for multiple modifications in single strand and/or hyphen (-) for next strand. For example ‘F,,T-T’, here ‘F’ and ‘T’(former one) are modifications for first strand as they are separated by double commas (,,) while the last ‘T’ is for second strand as it is separated by hyphen (-). The 21 modifications with single letter symbol are also provided in the form of dropdown list on the page for quick view of the user. Last alternative is for nucleotide positions for which the modifications need to be made (e.g. 2,5,7,10,11,12,13,14,15,16,,20,21-20,21). This option follows single comma (,) separator for same modification on multiple positions as 2,5,7,10,11,12,13,14,15,16 positional nucleotides will be modified with ‘F’ on first strand and ‘T’ modification will be performed on the 20 and 21 nucleotide positions on first strand. Likewise, ‘T’ (second one separated by hyphen) will be modified on 20 and 21 position of second strand.

Afterwards, the user can submit the query and will be directed to next page with input sequence, modifications, positions and output sequence (GFAGFAFGAFFFFFFFAA GTT-CUUGAAGAAGUCGUGCUGCTT) with modified nucleotides for confirmation. Download option for all this information is also provided on the page. Next user have to click on the modified sequence and will be directed the next page with output of this sequence, its length, toxicity derived by three SVM based toxicity models and knockdown efficacy [18].

### TOXsi-MODELS

TOXsi-Models provide the user with the choice to input the modified sequence both sense and antisense strands separated by hyphen as single string. This algorithm is based on the best three models of SVM based machine learning for predicting its toxicity. Therefore, upon submitting the query, user is directed to new page with the sequence, length and toxicity score values from the three best models and the knockdown efficacy in percentage.

### siOff-Tar

The siOff-Tar programme helps the user to screen the off target of the normal siRNA sequence against various model organisms namely: “Humans (*Homo sapiens*), Mouse (*Mus musculus*), Zebrafish (*Danio rerio*), Fruit fly (*Drosophila melanogaster*), Nematode worm (*Caenorhabditis elegans*)” [21].

Besides, help pages etc. are provided for helping the user to navigate and use the resource.

## Discussion

For improving RNAi technology, chemical modifications are used on the siRNA molecules, which can have toxicity effects on cells [4, 11]. Before making these chemical modifications it becomes essential to look for these effects as well as off potential targets of these siRNAs to minimize the harmful effects. Both these problems are addressed computationally with this web server. The importance of the web server lies in the fact that it is difficult to test these effects (especially toxicity due to different chemistries) experimentally particularly with different permutations or combinations of multiple chemical modifications. For example, for just five different modifications for 21 positions on one siRNA, we can have 210 (5*21*2) siRNAs to test [18]. Therefore, to avoid the heavy expenditure of resources and time for experimentally testing such chemical modified siRNA, computational algorithms can be of very much use.

Our results based on four different machine-learning techniques (SVM, LR, KNN and ANN) implied that different features performed differently for model development in terms of PCC values **(Table 2)**. Nearly on a particular feature all the different techniques performed more or less equally with certain variations. Nonetheless, the SVM based models performed best among all of them and were chosen for web implementation. Although, the mononucleotide composition (model-I) performed best but we added other two models (model-II and III) also as they included the positional information in addition to compositional information in the former model. The model-II involved the positional information of the antisense strand and model-III includes that of sense strand. The positional information or the pattern of the chemical modification is also very important in defining the knockdown effect of siRNA [22]. Furthermore, the independent validation of these three models also performed well as depicted by their scatter plot (**Figure 1**). Very good association is seen in all the three models between the predicted and actual knockdown scores with some outliers.

**Figure 1.**
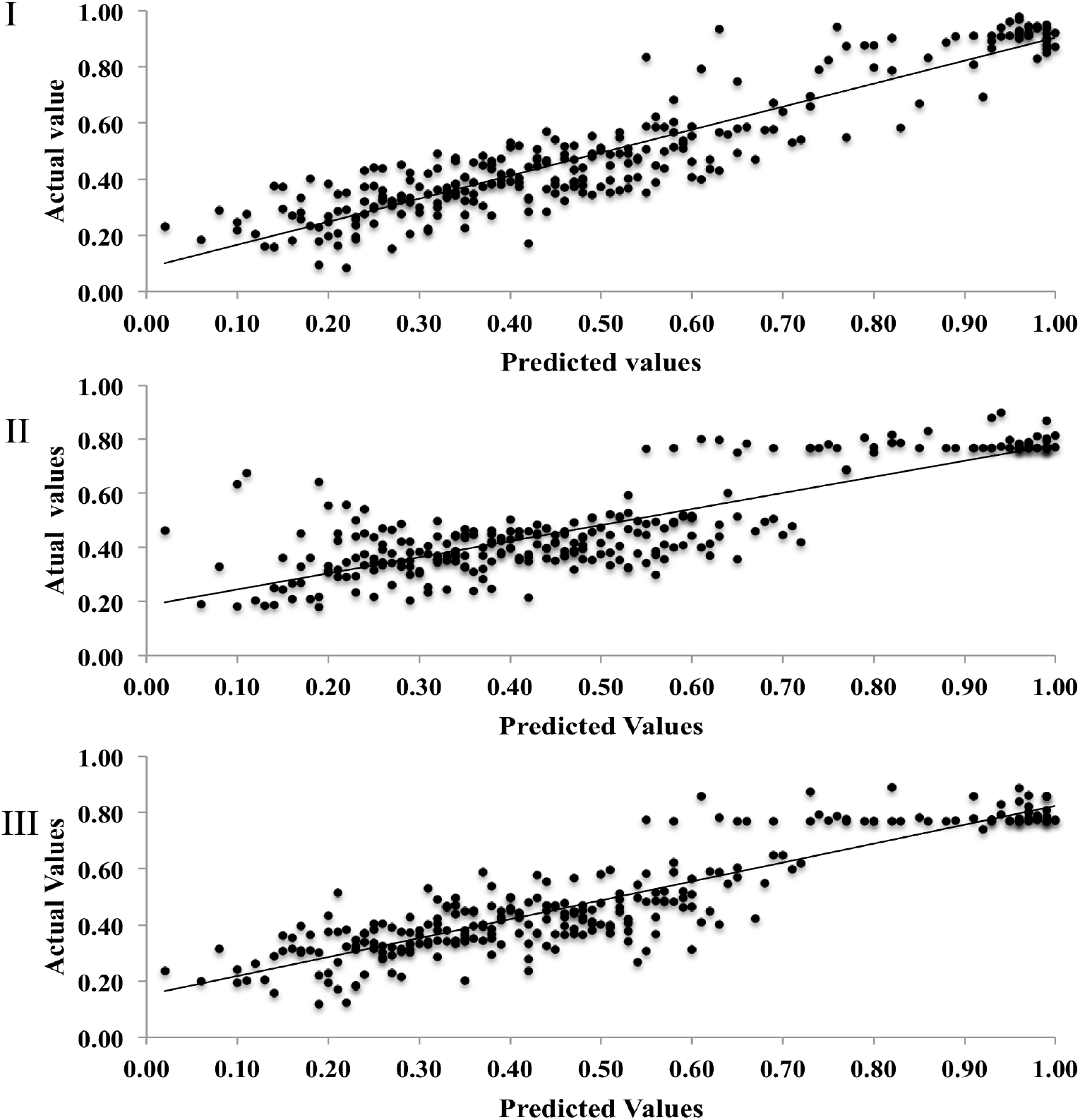
The Scatter plot of the three SVM based models that are integrated in the web server that shows very good positive correlation between the predicted and actual values: Here (I) is Mononucleotide composition, (II) is Mononucleotide composition with binary information of 24 nucleotides on antisense strand and (III) is Mononucleotide composition with binary information of 24 nucleotides on sense strand.

The siRNAs have the off-targets, similar to their miRNAs counterparts that can also be reduced by chemical modifications [14]. However, to be on safer-side we need to choose the siRNA with least number of off-targets, for which we have incorporated ‘siOff-Tar option’ in the web server. It provides an insight into the number of off targets in the genomes of five model organisms as per need of the user based on sequence similarity. Also other tools and web pages are provided for how to use guidelines as well as other used by the users.

## Conclusion

We provide TOXsiRNA as the first online web server to predict the toxicity of the chemically modified siRNAs (most common 21 modifications) in addition to predicting normal siRNAs as well as their off-targets. Its direct application is in the development of next generation siRNAs that involve chemical modifications or their combinations for potential therapeutics development as well as in research use. This web server is based on limited number of chemical modifications (only 21), which are most frequently used ones, and there is not much data for other modifications to develop their machine learning models. In future, we will try to improve the web server by incorporating other chemical moieties and related information in machine learning and model development for the same. We hope this web server will be useful to all the scientific community especially those working in siRNA based therapeutics development.

## Materials and Methods

### Data acquisition

We looked into the literature and checked the entire data for toxicity of chemically modified siRNAs. We narrowed down to 2749 siRNAs that were chemically modified with the values of the cellular toxicity available [4, 14]. In total 21 different chemical modifications (**Supplementary Table 1**) were decorated in these sequences in multiple repetitions and arrangements. We excluded the other data of modified siRNAs either because of less availability of data or their toxicity information was not available. Subsequently, the entire data set was normalized for range of values 0 to 1, where 0 was given the least toxicity score and 1 being the most toxic siRNA molecule (Microsoft Excel was employed for normalization).

### Feature selection of chemically modified siRNAs

We subsequently prepared the whole dataset of 2749 chemically modified siRNAs (A-2749) for the machine learning and divided the data into two sets one for training/testing and one for independent validation (with approximate ratio of 9:1) as T-2474 and IV-275 respectively [18]. Feature selection was performed on these sequences which was based on the mononucleotide composition, dinucleotide composition, binary pattern and their hybrids [23].

### Machine learning model development with Cross and Independent Validation

For machine learning we employed four techniques namely Support Vector Machines (SVM), Artificial Neural Networks (ANN), k-Nearest Neighbors (KNN) and Linear Regression (LR) [23]. For SVM, we engaged SVMlight package and for other three we used “Waikato Environment for Knowledge Analysis” (WEKA) package [20, 24, 25]. Subsequently, we used the T-2474 dataset for the 10-fold cross validation using all the above techniques [17]. In this method, the data is divided into ten equal datasets and nine are used for model development and one dataset is used for its validation [23]. The cycle is repeated ten times and each time the validations dataset is different. In independent validation, the dataset that is used for validation is not used for model development and is independent of model development, in contrast to cross validation [17]. We used two methods for independent validation of models namely ‘systematic’ and ‘random’. In the former process, we divide the data sequentially into training and independent validation datasets (one-tenth of overall) based on sequential increase or decrease of the data values (toxicity values). For example after aligning the entire data ascending order of toxicity values, we choose every n^th^ sample (where n is a number say 2^nd^, 5^th^, 8^th^) after each ten sequences [23]. This set is used for independent-validations, whereas rest of the dataset is used for training testing of that model. For random dataset based model validation, we randomly select a one-tenth datasets, exclude it for model development and use it at last for validating the model independently [17].

### Other tools

For designing of the normal siRNAs we have integrated the SVM based models from our earlier work describes elsewhere [18]. Furthermore, for identification of the off targets, we have integrated a tool based on BLAST (Basic Local Alignment Search Tool) that reveals the off-targets of the unmodified siRNAs. Five different model organisms (Homo sapiens, Mus musculs, Danio rerio, Drosophila melanogaster, Caenorhabditis elegans) are provided to choose from according to the users choice [26]. Tools for designing and creating the chemically modified siRNA sequences etc. are also integrated in the web server based on in-house scripts [18]. Hyper Text Markup Language (HTML), Cascading Style Sheets (CSS), Hypertext Preprocessor (PHP), JavaScript (JS), Perl and Python were employed to develop the web server [27].

## Supporting information

Graphical Abstract

## Supplementary data

Supplementary file:

## Acknowledgment

Supported by Council of Scientific & Industrial Research (CSIR) Institute of microbial technology (IMTECH) (OLP143) and Department of biotechnology (DBT) GAP0001

## Conflict of interest

Declared none

**Database URL**: http://bioinfo.imtech.res.in/manojk/toxsirna

