## Supplementary figures and images for "TOXsiRNA: A web server to predict the toxicity of chemically modified siRNAs"

### Graphical Abstract

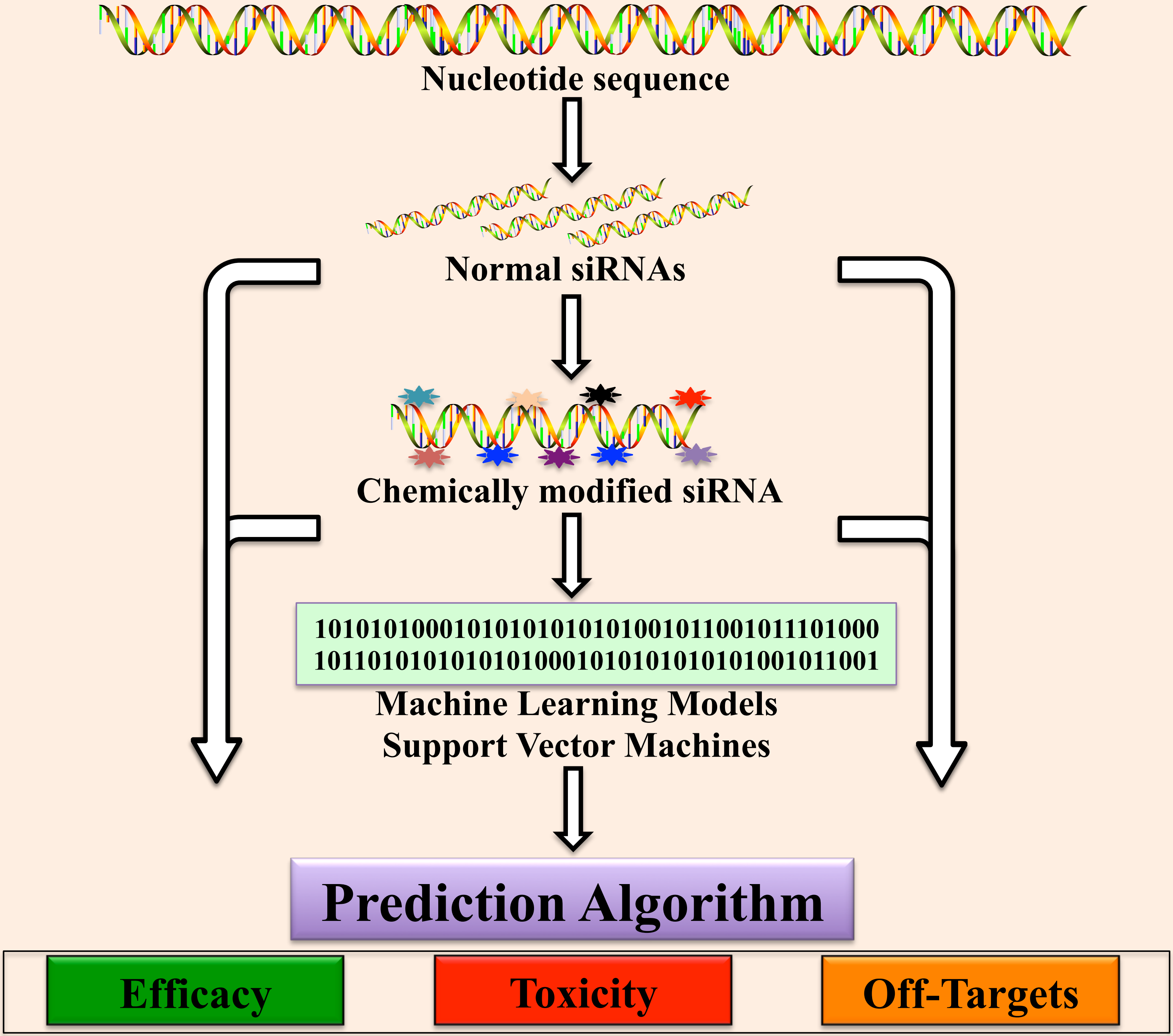
