## Supplementary File for "TOXsiRNA: A web server to predict the toxicity of chemically modified siRNAs"

Showkat Ahmad Dar

Virology Discovery Unit and Bioinformatics Centre,

Institute of Microbial Technology, Council of Scientific and Industrial Research, Sector 39A, Chandigarh-160036, India

Manoj Kumar

Virology Discovery Unit and Bioinformatics Centre,

Institute of Microbial Technology, Council of Scientific and Industrial Research, Sector 39A, Chandigarh-160036, India

**Supplementary file:**

**Supplementary Table 1.** The list of chemical modifications used with their single letter codes.

| **Serial No.** | **Single letter code** | **Chemical modification** |
| --- | --- | --- |
| **1** | D | 2'-Aminoethoxymethyl Nucleic Acid |
| **2** | E | 2'-Aminoethyl Nucleic Acid |
| **3** | F | 2'-Fluoro Nucleic Acid |
| **4** | H | 4'-C-Hydroxymethyl Deoxyribonucleic acid |
| **5** | I | Altritol Nucleic Acid |
| **6** | J | 2'-Aminopropyl Nucleic Acid |
| **7** | K | 2'-aminopropoxymethyl Nucleic Acid |
| **8** | L | Locked Nucleic Acid |
| **9** | M | 2'-Deoxy-2'-N,4'-C-Ethylene-Locked Nucleic Acid |
| **10** | N | Alfa-L-Locked Nucleic Acid |
| **11** | O | 2'-Cyanoethyl Nucleic Acid |
| **12** | P | 2',4'-Carbocyclic-Ethylene-bridged Nucleic Acid-Locked Nucleic Acid |
| **13** | Q | 2',4'-Carbocyclic-Locked Nucleic Acid-Locked Nucleic Acid |
| **14** | R | 2'-Guanidinoethyl Nucleic Acid |
| **15** | S | Hexitol Nucleic Acid |
| **16** | V | Oxetane-Locked Nucleic Acid |
| **17** | W | 2'-N-Pyren-1-yl Methyl-2'-Amino-Locked Nucleic Acid |
| **18** | X | Unlocked Nucleic Acid |
| **19** | Y | 2'-O-Methyl Nucleic Acid |
| **20** | Z | 2'-Deoxy Nucleic Acid |
| **21** | b | 2'-Deoxy-2'-Fluoro Nucleic Acid |

Normal nucleotides (A, T, G, C, U) are not shown in the table.
